# The eukaryotic replisome requires an additional helicase to disarm dormant replication origins

**DOI:** 10.1101/2020.09.17.301366

**Authors:** Jake Hill, Patrik Eickhoff, Lucy S. Drury, Alessandro Costa, John F.X. Diffley

## Abstract

Origins of eukaryotic DNA replication are ‘licensed’ during G1 phase of the cell cycle by loading the six related minichromosome maintenance (MCM) proteins into a double hexameric ring around double-stranded DNA. In S phase, some double hexamers (MCM DHs) are converted into active CMG (Cdc45-MCM-GINS) helicases which nucleate assembly of bidirectional replication forks. The remaining unfired MCM DHs act as ‘dormant’ origins to provide backup replisomes in the event of replication fork stalling. The fate of unfired MCM DHs during replication is unknown. Here we show that active replisomes cannot remove unfired MCM DHs. Instead, they are pushed ahead of the replisome where they prevent fork convergence during replication termination and replisome progression through nucleosomes. Pif1 helicase, together with the replisome, can remove unfired MCM DHs specifically from replicating DNA, allowing efficient replication and termination. Our results provide an explanation for how excess replication license is removed during S phase.

## Introduction

Eukaryotes initiate genome duplication from large numbers of sites, known as DNA replication origins, distributed along multiple chromosomes^1^. At origins, the six minichromosome maintenance (MCM) proteins are loaded into head-to-head double heterohexamers (MCM DH) by the Origin Recognition Complex (ORC), Cdc6 and Cdt1. Each MCM DH can then be converted into two divergent replication forks in which the MCM hexamers act as the motor of the CMG replicative helicase.

To ensure replication origins fire just once in each cell cycle, MCM loading (licensing) is restricted to the G1 phase of the cell cycle. Thus, once DNA replication begins in S phase, MCM can no longer be loaded. It is therefore crucial that sufficient MCM is loaded during G1 phase to replicate the entire genome during S phase. This is accomplished by loading an excess of MCM DH onto chromosomes. A subset of these MCM DHs are converted into replication forks during S phase whilst the remaining MCM DHs serve as backups known as ‘dormant’ origins^2, 3, 4^. Though normally inactive, dormant origins can be activated if passive replication through them is prevented, for example by stalling of the replication fork from a neighbouring origin^3, 4^. Consequently, dormant origins are especially important in promoting complete replication during replication stress^5, 6, 7^.

The MCM DH in yeast is an extraordinarily stable structure, resistant to 2M NaCl^8^. But it is also a dynamic structure: because it associates as a ring around double-stranded DNA^9, 10^, it is free to ‘slide’ and can be pushed to distant sites by DNA translocases like the CMG helicase^11^ and RNA polymerase where it can act to initiate replication^12^.

The original licensing factor model posited that the act of replication in some way ‘erased’ the license from DNA as it is replicated^13, 14, 15^. It may be that the replisome itself in some way removes unused MCM DH from chromatin. Alternatively, additional non-replisome factors may be involved. Two candidates have emerged from analysis of yeast mutants: Mcm10 and Rrm3. Hypomorphic *mcm10* mutants^16^ and *rrm3Δ* cells^17^ both display more significant ‘pauses’ at dormant replication origins *in vivo* than wild type cells by two-dimensional gel electrophoresis analysis of replication intermediates. This implies replisomes have difficulties traversing these sequences in the absence of Mcm10 and/or Rrm3. It remains unclear whether replisome pausing was caused by some aspect of the origin DNA sequence, the binding of ORC, the binding of some other factor, like Abf1, or the presence of the MCM DH.

## Results

### Okazaki fragment maturation

We have previously reconstituted regulated DNA replication of naked DNA and chromatin templates with purified yeast proteins^10, 18, 19, 20^. Replication in this system proceeds at rates similar to those seen *in vivo* and generates equal amounts of leading and lagging strand products^18, 20^. During the process of Okazaki fragment maturation, RNA primers from each Okazaki fragment must be removed and Okazaki fragments then ligated together; two flap endonucleases, Fen1 and Dna2, along with the DNA ligase, Lig1, have been implicated in this process^21^. Addition of purified Fen1, Dna2 and Lig1 (Fig. 1A) during replication of a naked DNA template (Fig. 1B) led to disappearance of Okazaki fragments and conversion of the DDK-dependent leading and lagging strand products (Fig. 1C lanes 2 and 3) to higher molecular weight (MW) products on denaturing alkaline agarose gels (Fig. 1C lanes 4 and 5). Generation of high MW products required Fen1 but not Dna2 (Fig. 1A lanes 8-11). In the absence of Lig1, addition of Fen1 caused accumulation of high molecular weight labelled product even in the absence of DDK, indicating that this product is not the result of replication, but some non-specific synthesis (Fig. 1C lanes 6 and 7). Thus, consistent with previous work^22^, Fen1 and Lig1 are necessary and sufficient to process and ligate Okazaki fragments *in vitro*. Dna2 increased the size distribution of Okazaki fragments (Fig. 1C, lanes 8 and 9), but did not generate high MW products indicating that it cannot fully substitute for Fen1. Synthesis of each Okazaki fragment is initiated by DNA polymerase α, but completed by DNA polymerase δ with its processivity factor, PCNA. Fig. 1D shows that generation of high MW products by Fen1 and Lig1 also required DNA polymerase δ and PCNA indicating that complete Okazaki fragment synthesis of required for maturation. Replication of chromatinised templates requires the histone chaperone FACT and its co-factor, Nhp6 (Ref ^18^ and Fig. 1E lanes 1 and 2). Fen1 and Lig1 were also necessary and sufficient for generation of high MW products on chromatin templates (Fig. 1E). Taken together, our results show that Fen1 and Lig1, along with complete Okazaki fragment synthesis, are required for Okazaki fragment maturation.

**Figure 1.**
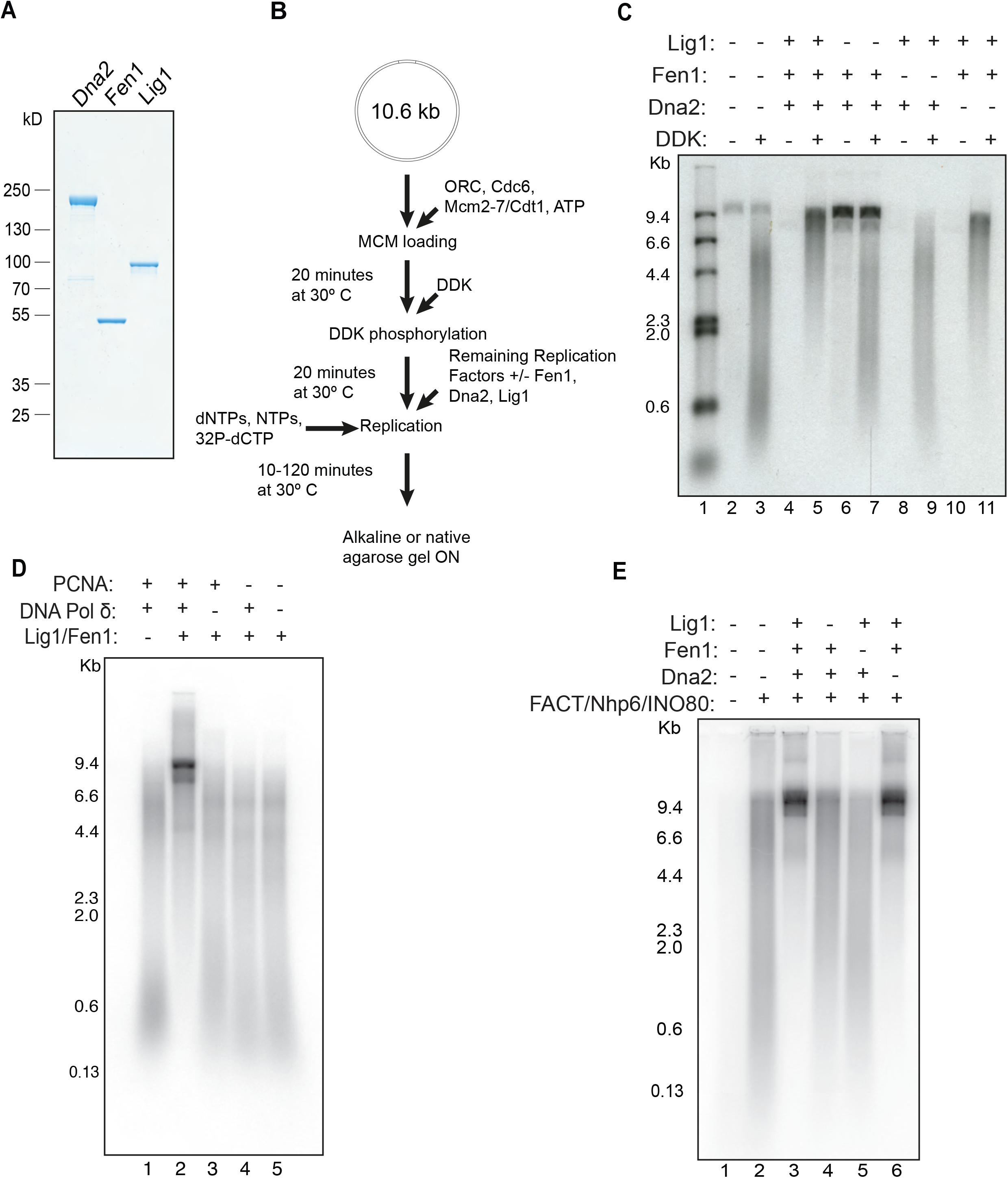
Okazaki fragment maturation. **a**, Purified Okazaki fragment maturation proteins analysed by SDS-PAGE with Coomassie staining. **b**, General scheme for replication reactions. **c**, Reactions performed as in **b** with various combinations of Dna2, Fen1, and Lig1. Products were separated on an alkaline agarose gel. **d**, Reactions performed with various combinations of Fen1, Lig1, PCNA, and DNA Polymerase δ. Products were separated on an alkaline agarose gel. **e**, Reactions performed as in **c** on chromatinised DNA.

### Termination of DNA replication in vitro

The high MW products we saw in alkaline agarose gels from reactions containing Fen1 and Lig1 were roughly the size of the template plasmid (10.8 kb) and twice the size of the leading strand products (∼5kb). This suggested that Okazaki fragments were being ligated not only to each other, but also to the leading strand products from the divergent replication fork, either at the origin, at the terminus, or both. We, therefore, wanted to ascertain whether complete replication was occurring in our reactions. To test this, we ran replication products on neutral agarose gels. Fig. 2A shows that bulk DNA, visualised by Ethidium Bromide (EtBr) staining after electrophoresis, coincided with form II/I^0^ DNA (nicked/relaxed closed circular^23^) with a cluster of topoisomers below. This happened with or without DNA replication (+ or - DDK) because topoisomerase in the reactions relaxed all of the input supercoiled plasmid DNA. As shown in the autoradiogram (compare Autoradiogram, lanes 1 and 2), there were two roughly equal populations of DDK-dependent replicated products. One population migrated more slowly than any forms of the monomeric plasmid; these products presumably represent incomplete replication products, or replication intermediates (RIs). The second population of products appeared identical to the EtBr stained gel corresponding to the relaxed plasmid with topoisomers. Digestion of replication products with the restriction endonuclease XhoI, which has a single cleavage site in the template plasmid, produced a single major band of ∼10.8kb corresponding to Form III (linear) of the plasmid (Fig. 2A lane 3). The slow running replication intermediates (RIs) were also converted to a faster migrating series of bands after linearization. Sup. Fig. 1A shows that a full-length nascent strand can be seen after XhoI digestion by alkaline gel electrophoresis as well. Neutral-alkaline two-dimensional gel analysis of replication products after XhoI digestion (Supp. Fig.2B) showed that RIs resolve into a ‘C’-shaped arc in the second dimension. This is consistent with them comprising a set of X-shaped molecules that have not terminated replication, with the slowest migrating forms in the first dimension corresponding to molecules with equal length arms^24^ (Supp. Fig. 2C). From these experiments we conclude that approximately half of the products are fully replicated covalently closed circular DNA which have terminated replication whilst the other half are late RIs that have not terminated replication. Although termination was efficient, it was slow relative to replication (Fig. 2B, C): bulk replication was complete by 10-20 minutes, but termination continued over the course of the experiment, reaching almost 60% after two hours (Fig. 2C).

**Figure 2.**
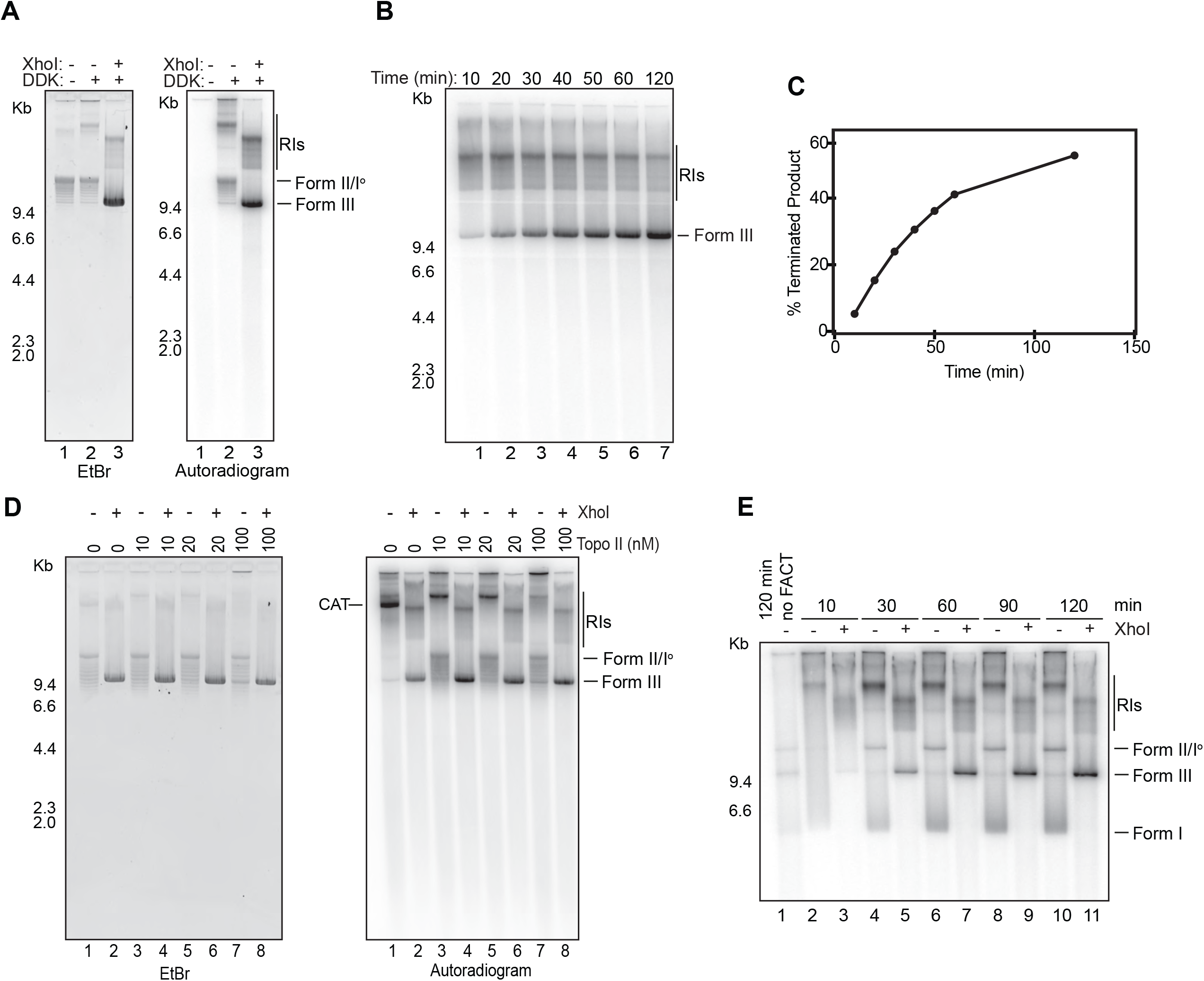
Termination of DNA replication in vitro. Unless otherwise stated, all subsequent reactions were performed as in Fig. 1b. **a**, Following replication half of the +DDK products were linearised with XhoI. They were then separated on a native agarose gel and stained with ethidium bromide (left) prior to drying and subjecting to autoradiography (right). **b**, Time course of a reaction performed as in **a**. The products were linearised following replication and separated on a native agarose gel. **c**, Quantification of the terminated (linear) product as a percentage of the total product in each lane of **b. d**, Titration of Topoisomerase II. **e**, Time course of replication on chromatin. Products were split and half were linearised prior to separation on a native agarose gel.

Replication of covalently closed circular plasmids requires topoisomerase activity, and either Top1 or Top2 can provide this activity during replication *in vivo* and *in vitro*^19, 25^. Supp. Fig. 2 shows that Top2 can support complete replication including termination in the absence of Top1 and the addition of Top1 had little effect on replication products (Fig. 2D and Supp. Fig. 2). Top1 also supported efficient replication in the absence of Top2 (Fig. 2D, lanes 1,2). Monomeric circles were missing from a reaction lacking Top2 (lane 1), but instead a novel high MW band corresponding to catenated dimers (Cat) was present. Monomeric linear molecules appeared after XhoI digestion (lane 2) indicating that termination had occurred, but decatenation had not. These results indicate that Top2 is required for decatenation, but is not essential for termination. We note there was a small decrease in the amount of Form III product in the absence of Top2 (Compare Fig. 2D lanes 2 and 4) suggesting Top2 may play a non-essential but stimulatory role in termination.

We next examined replication on chromatin templates by neutral agarose gel electrophoresis. The appearance of Form III products that accumulate over two hours after linearisation shows that termination occurred with similar efficiency on chromatinised templates. In contrast to replication of naked DNA, monomeric products from reactions lacking XhoI (even number lanes) were predominately supercoiled Form I instead of relaxed Form I^0^. This suggests that parental nucleosomes are being redeposited onto nascent DNA during replication, which is consistent with our previous work showing nascent DNA after replication of chromatin displays the classic micrococcal nuclease digestion pattern of nucleosomal arrays^18^. This provides further evidence that histones evicted ahead of the replication fork are efficiently transferred to nascent DNA behind the fork *in vitro*. Taken together, our conditions support replication termination on naked DNA and chromatin without requiring additional factor(s)^26^.

### The replisome cannot displace MCM double hexamers

We previously found that active CMG can push MCM DHs, which are topologically bound around double stranded DNA, against a covalent protein-DNA roadblock, generating ‘trains’ of double hexamers^11^. The presence of these trains suggested that CMG alone cannot displace MCM DHs. Single molecule studies have shown that CMG is a relatively slow helicase which frequently stalls and backtracks^27^ and, thus, may not be able to generate sufficient force to displace MCM DH from DNA. We therefore wondered whether the complete replisome^20^, could displace excess double hexamers from DNA. To address this, we asked whether DNA replication was affected by excess MCM loading. To accomplish this, we increased MCM loading by increasing the amount of MCM•Cdt1 in our loading reactions^11^. This was done under conditions where MCM loading does not require a replication origin and therefore occurs randomly on the plasmid template^10^. As shown in Fig. 3A, the amount of terminated product (Form III) after replication decreases with increasing amounts of MCM whilst the amount of RI increases. The appearance of terminated product was not inhibited when excess soluble double hexamers were added at the onset of replication, when their loading is prevented by CDK (Supp. Fig. 3A). This suggests that excess MCM specifically blocks termination. To investigate this further, we examined replication products by electron microscopy after negative staining. In addition to smaller complexes — mostly MCM single and double hexamers — we saw examples of long filamentous structures (Supp. Fig. 3B, C). These were only seen in reactions with excess loaded MCM. Fig. 3C shows an example of such a structure. Closer examination reveals that these filamentous structures comprise multiple double hexamers (Fig. 3e, blue panels); 48% of these structures were capped at one end by a CMG whilst 31.2% were sandwiched between two CMGs (Fig. 3e, green panels and Supp. Table 1). Other components of the replisome were visible, including DNA polymerase ε and Ctf4 (Fig. 3e, green panels). Their position on CMG, together with previous work on the orientation of the CMG helicase at the fork^11, 28, 29^ unambiguously shows that these structures represent MCM double hexamers trapped between two converging CMG-containing forks. Together, these data indicate that inactive double hexamers are pushed ahead of the replisome, accumulate at converging forks, and block termination.

**Table 1.**
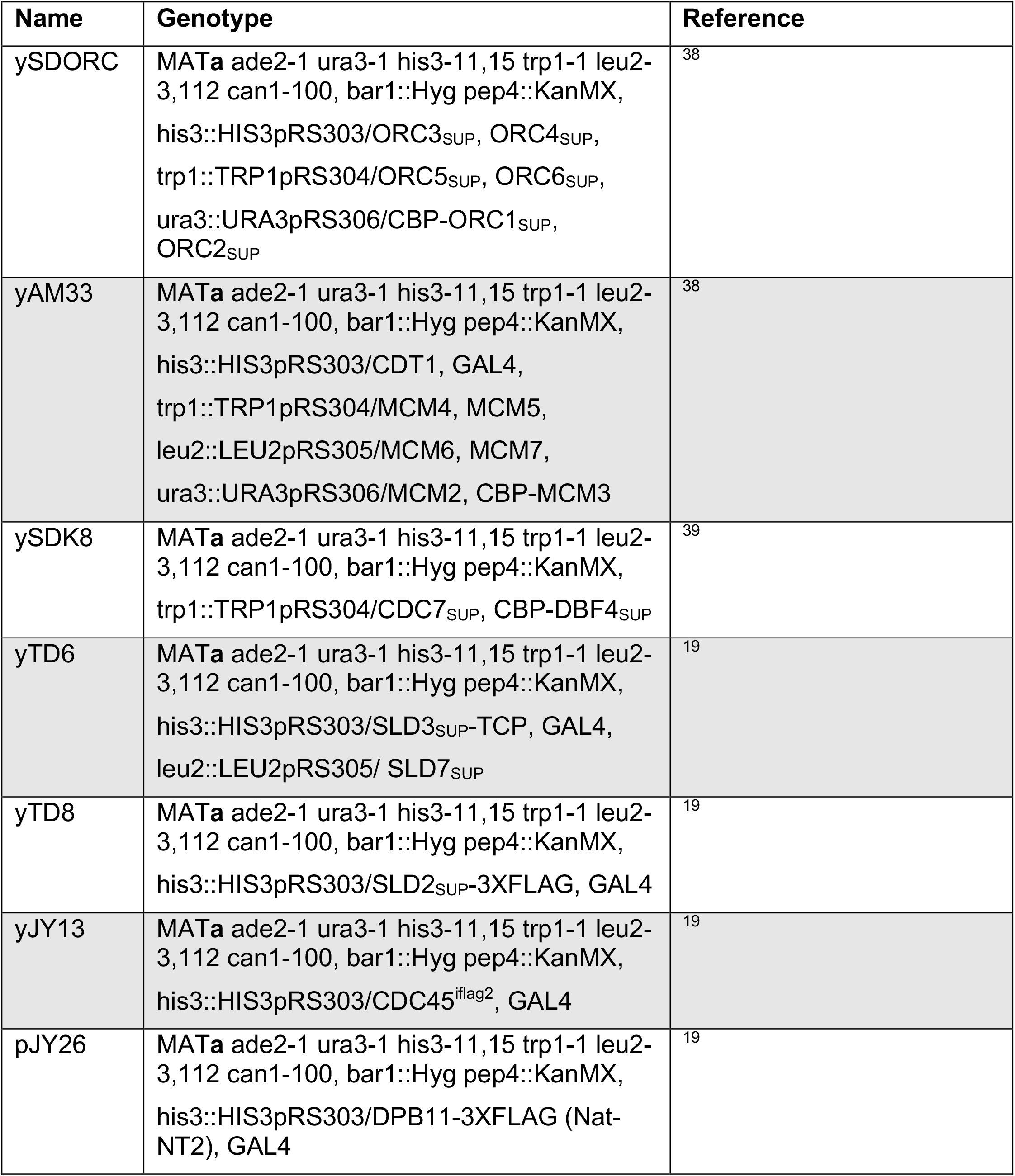
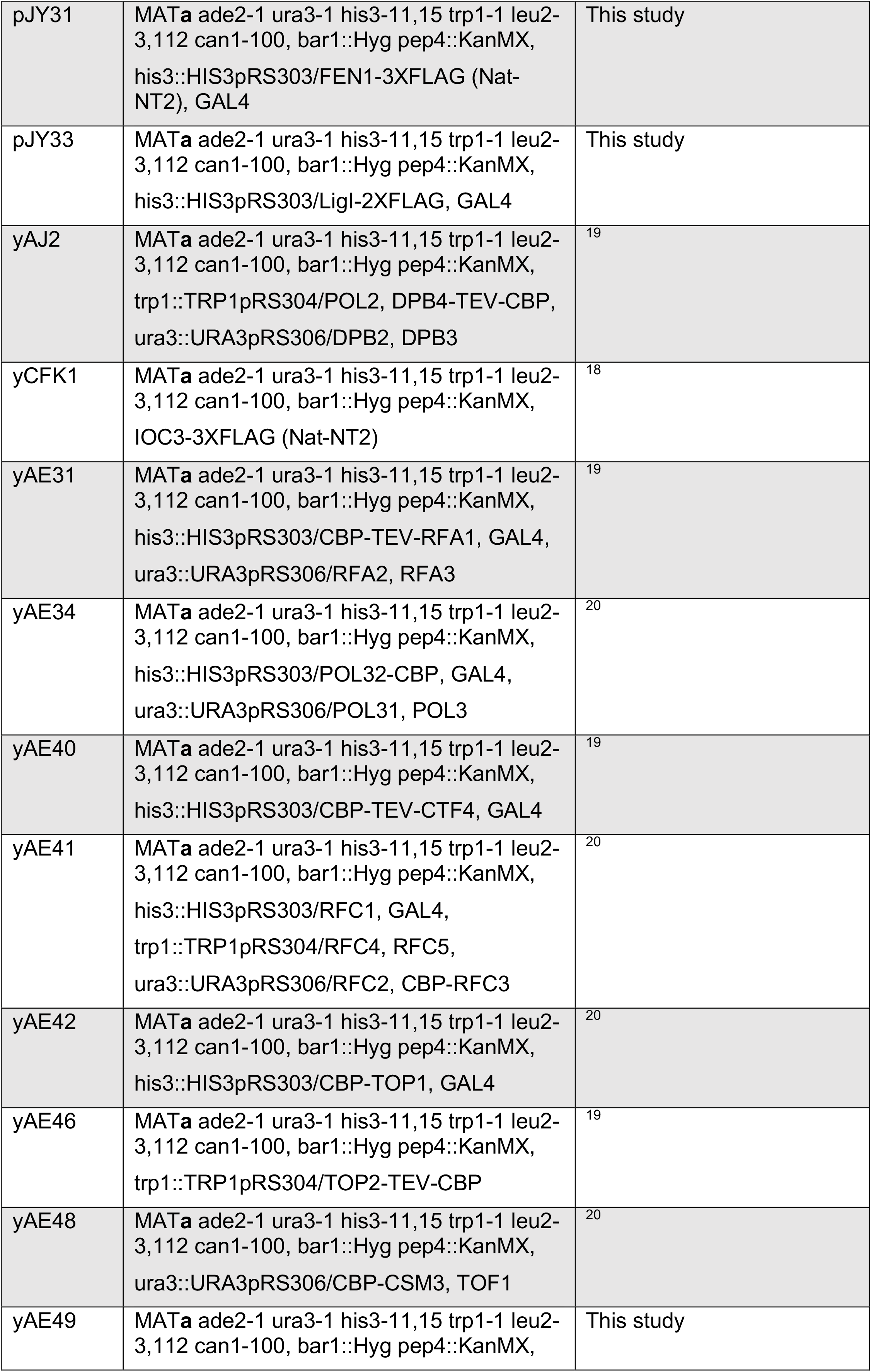
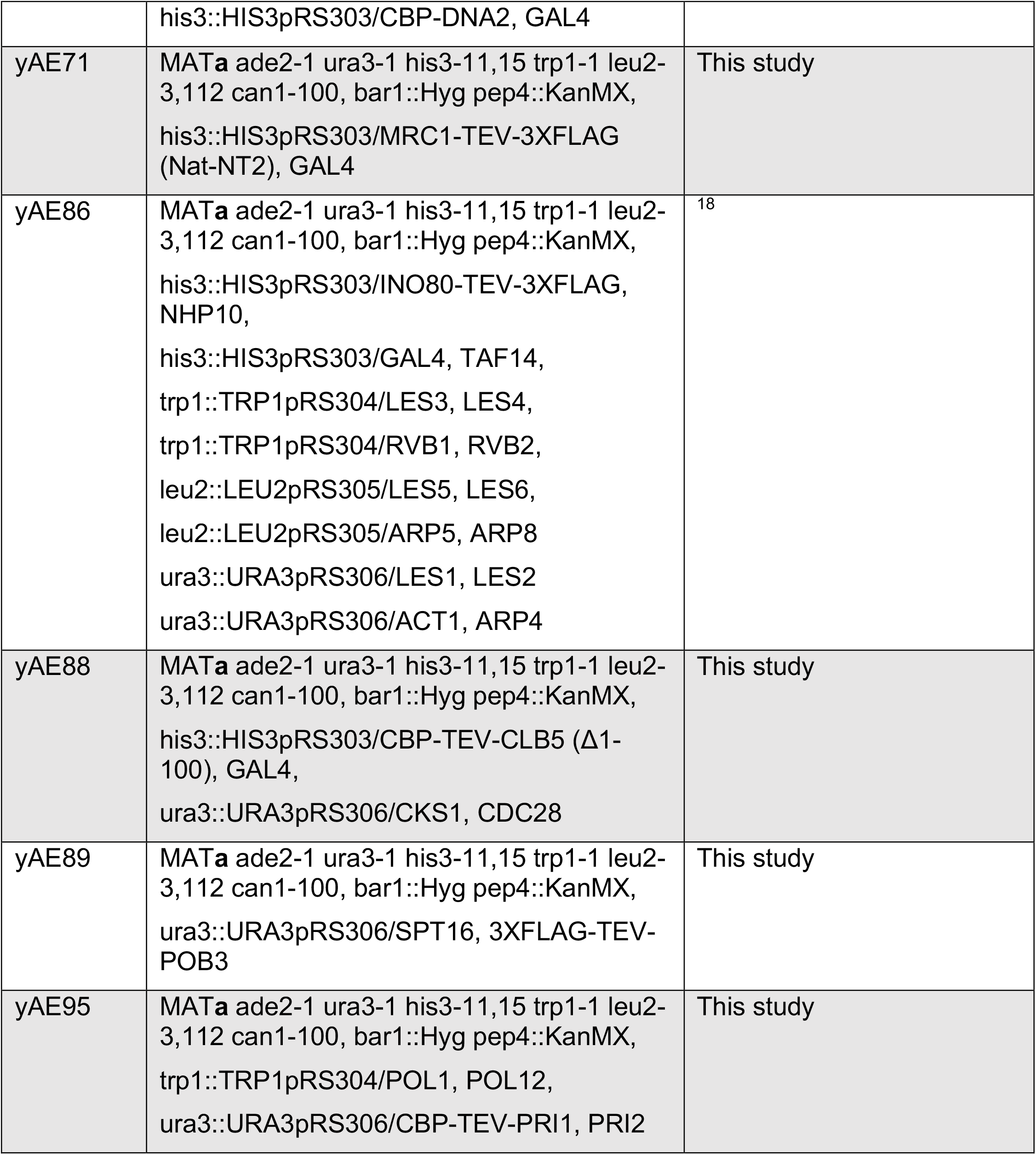
Yeast strains.

**Table 2.**
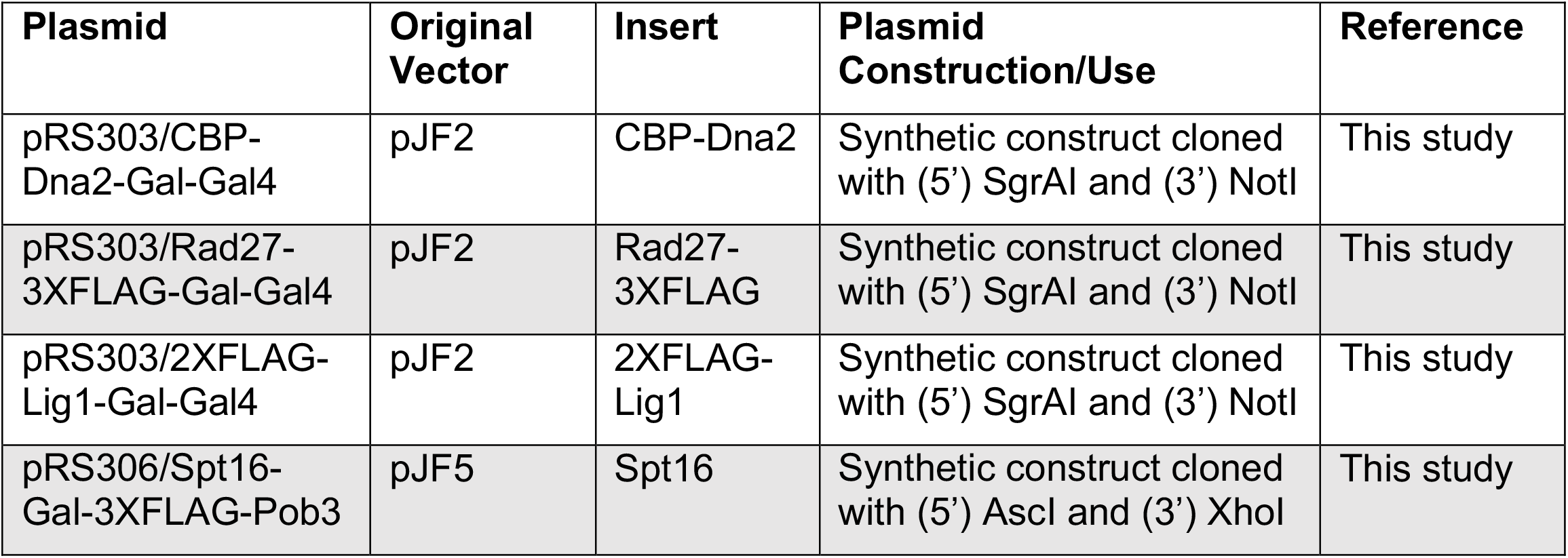
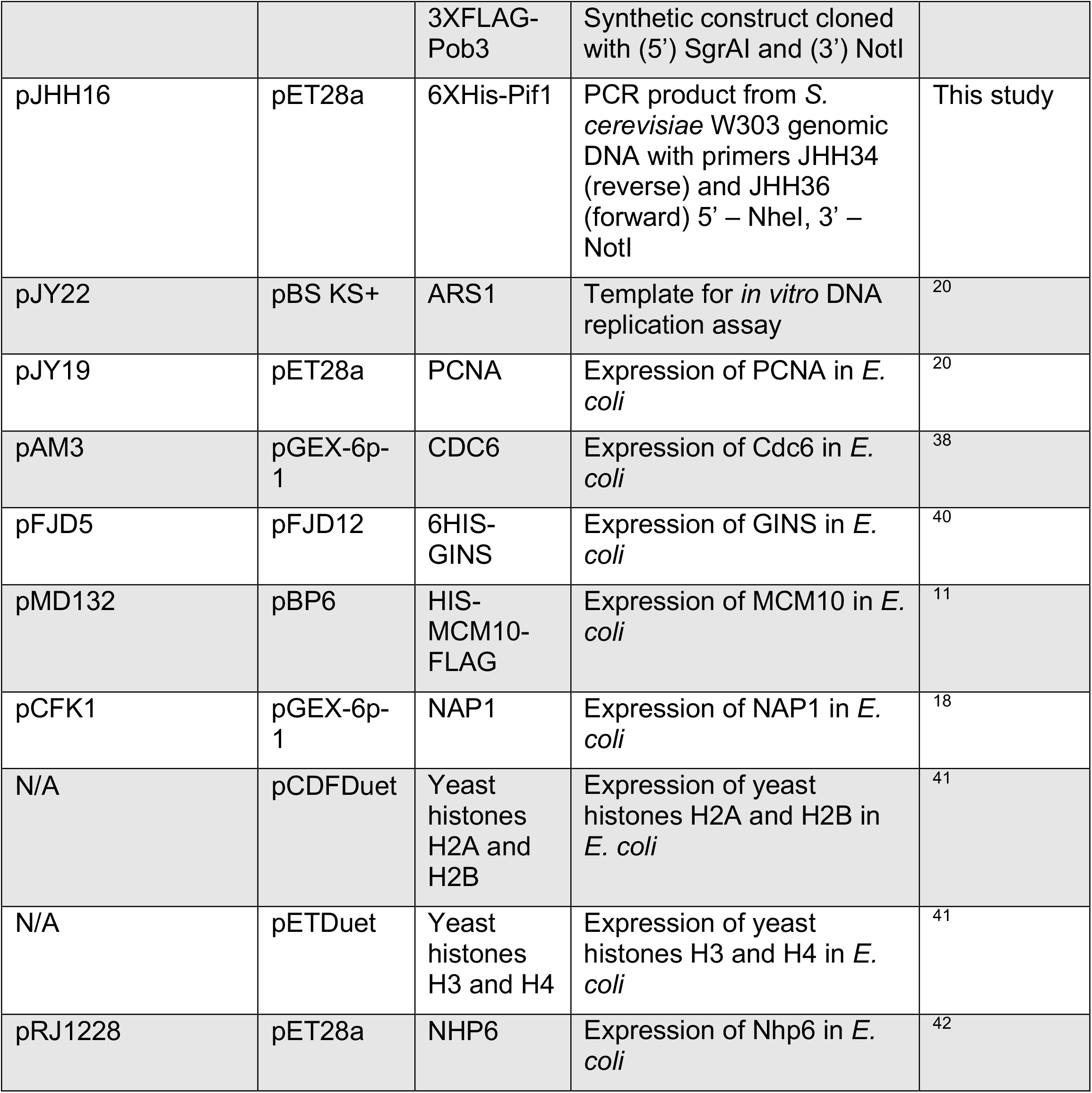
Plasmids.

| Plasmid | Original Vector | Insert | Plasmid Construction/Use | Reference |
| --- | --- | --- | --- | --- |
| pRS303/CBP-Dna2-Gal-Gal4 | pJF2 | CBP-Dna2 | Synthetic construct cloned with (5') SgrAI and (3') NotI | This study |
| pRS303/Rad27-3XFLAG-Gal-Gal4 | pJF2 | Rad27-3XFLAG | Synthetic construct cloned with (5') SgrAI and (3') NotI | This study |
| pRS303/2XFLAG-Lig1-Gal-Gal4 | pJF2 | 2XFLAG-Lig1 | Synthetic construct cloned with (5') SgrAI and (3') NotI | This study |
| pRS306/Spt16-Gal-3XFLAG-Pob3 | pJF5 | Spt16 | Synthetic construct cloned with (5') AscI and (3') XhoI | This study |
|  |  | 3XFLAG-Pob3 | Synthetic construct cloned with (5') SgrAI and (3') NotI |  |
| pJHH16 | pET28a | 6XHis-Pif1 | PCR product from *S. cerevisiae* W303 genomic DNA with primers JHH34 (reverse) and JHH36 (forward) 5' – NheI, 3' – NotI | This study |
| pJY22 | pBS KS+ | ARS1 | Template for *in vitro* DNA replication assay | 20 |
| pJY19 | pET28a | PCNA | Expression of PCNA in *E. coli* | 20 |
| pAM3 | pGEX-6p-1 | CDC6 | Expression of Cdc6 in *E. coli* | 38 |
| pFJD5 | pFJD12 | 6HIS-GINS | Expression of GINS in *E. coli* | 40 |
| pMD132 | pBP6 | HIS-MCM10-FLAG | Expression of MCM10 in *E. coli* | 11 |
| pCFK1 | pGEX-6p-1 | NAP1 | Expression of NAP1 in *E. coli* | 18 |
| N/A | pCDFDuet | Yeast histones H2A and H2B | Expression of yeast histones H2A and H2B in *E. coli* | 41 |
| N/A | pETDuet | Yeast histones H3 and H4 | Expression of yeast histones H3 and H4 in *E. coli* | 41 |
| pRJ1228 | pET28a | NHP6 | Expression of Nhp6 in *E. coli* | 42 |

**Table 3.**
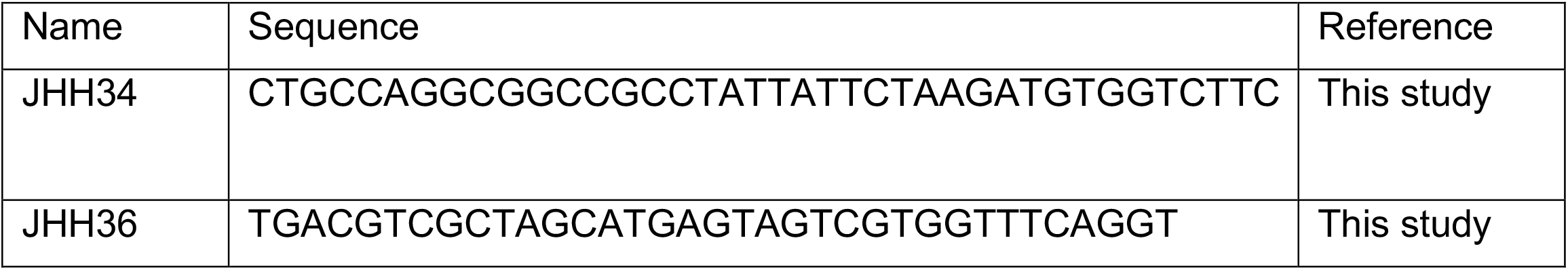
DNA Oligonucleotides.

**Figure 3.**
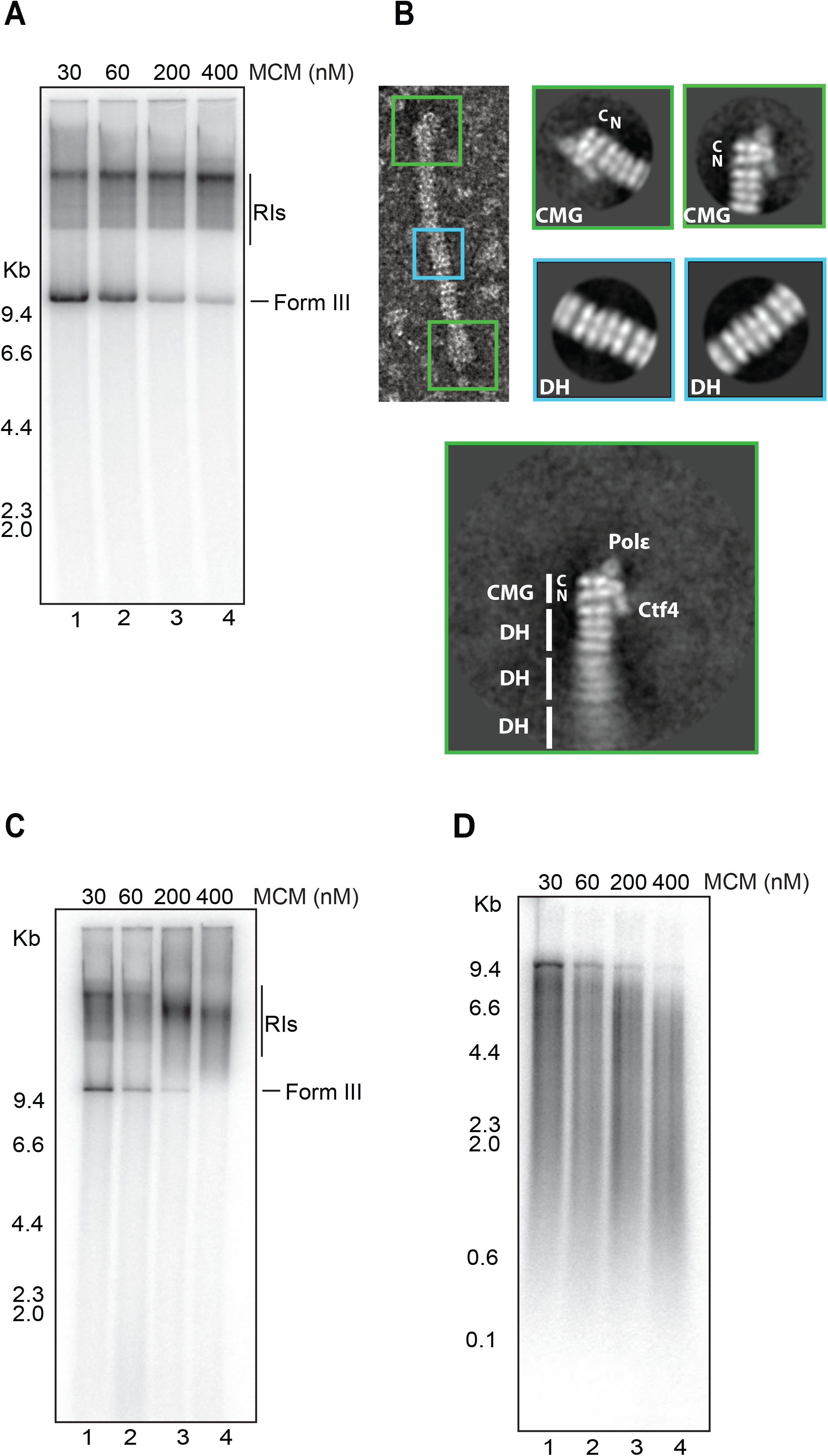
The replisome cannot displace MCM double hexamers. **a**, Reactions with increasing amounts of MCMs loaded onto DNA. Products were linearised and separated on a native agarose gel. **b**, Representative MCM train from replication of plasmid template DNA with an excess of loaded MCMs. 2D averages of train termini (green) and middle (blue) particles reveals MCMs between two converging CMGs **c, d** Reactions on chromatin with increasing amounts of loaded MCMs. Products were split in half and separated on a native agarose gel (**c**) or alkaline agarose gel (**d**).

To examine the effects of excess MCM DH on chromatin replication, we first loaded excess MCM randomly onto naked DNA, then chromatinised the plasmid and replicated it. Fig. 3C shows that excess MCM DHs also prevented accumulation of terminated products (Form III) on chromatinised templates in a dose-dependent manner. However, in contrast to reactions on naked DNA, the migration of RIs changed significantly with increasing MCM concentration, suggesting a more general inhibition of elongation. Indeed, when we analysed products on alkaline agarose gels, we found that the lengths of replication products decreased with increasing MCM concentration. Thus, excess double hexamers block replication of chromatin during elongation at a stage prior to termination, suggesting that the replisome is unable to push MCM DHs through nucleosomes.

### Pif1 can unload MCM double hexamers with the replisome

These results show that the replisome is unable to remove unfired MCM double hexamers indicating that an additional factor (or factors) is required to accomplish this. Because of the genetic evidence, we were interested in testing whether the Pif1 family of helicases could help remove MCM DHs. We were unable to produce sufficient amounts of active Rrm3, which has been directly implicated in replication through dormant origins^17^, but we were able to express and purify its relative, Pif1 (Fig. 4A). Previous work has shown that Pif1 can promote replication termination using purified proteins^26^. Consistent with this, we saw that addition of Pif1 increased the rate of termination, though we did not see an increase in the overall amount of terminated product at late time points (Supp. Fig. 4A,B). Pif1 also improved termination on chromatinised templates (Supp. Fig. 4C,D). Addition of Pif1 also generated a product that remained in the well after electrophoresis (‘W’ in Fig. 4B and Supp. Fig. 4A) along with a series of faint bands running faster than the terminated products (‘Low MW’ in Fig. 4B and Supp. Fig. 4A).

**Figure 4.**
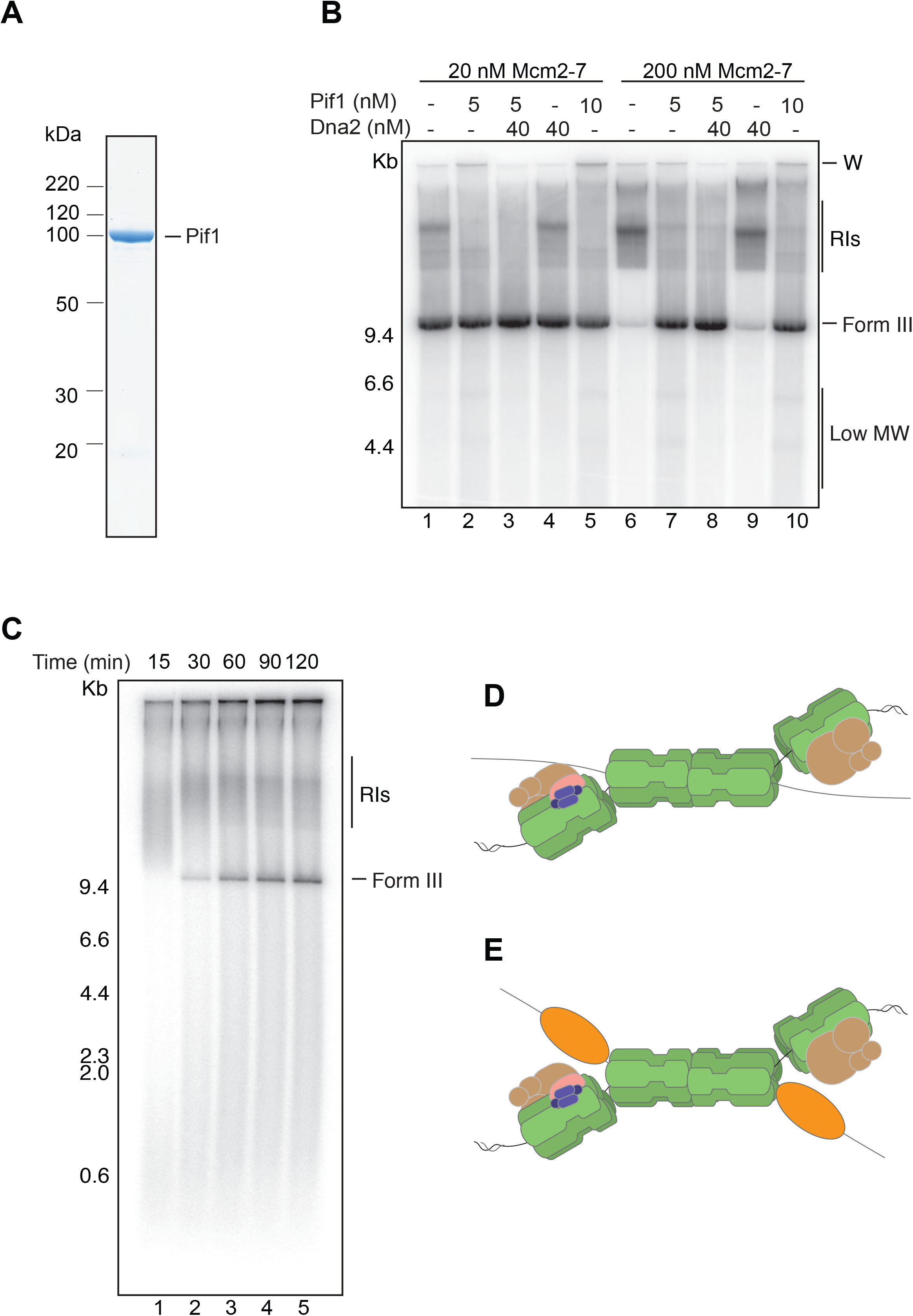
Pif1 can unload MCM double hexamers with the replisome. **a**, Purified Pif1 analysed by SDS-PAGE with Coomassie staining. **b**, Reactions performed with or without an excess of loaded MCMs in the presence or absence of Pif1 and/or Dna2. Products were linearised, separated on a native agarose gel, then dried and subjected to autoradiography. **c**, Time course of replication on chromatin with an excess of loaded MCMs in the presence of Pif1. Products were linearised and separated on a native agarose gel. **d, e** Schematics depicting two converging replication forks with and without Pif1, suggesting a possible role in removing unfired MCM DHs.

At low MCM concentrations (Fig. 4B lanes 1-5) addition of Pif1 did not change the amount of terminated product, but did reduce the amount of RIs (Fig. 4B lanes 1 and 2). In the presence of excess MCM, we saw, as before, a dramatic reduction in the amount of terminated product with a commensurate increase in the amount of RIs (Fig. 4B, compare lanes 1 and 6). Addition of Pif1 to this reaction completely restored the appearance of terminated product (Fig. 4B, compare lanes 6 and 7). We detected a significant drop in the number of ‘trains’ of MCM DHs in these reactions: whilst 1,700 trains per grid square were observed in the absence of Pif1, only 305 trains per grid square were seen when Pif1 was added. This difference was not due to variation in MCM particle concentration, as both samples contained an average of 400 particles per micrograph, not incorporated in trains. We found Dna2, another 5’—>3’ helicase, even at a much higher concentration, had no discernible effect on the products formed either with or without Pif1. We note that addition of Dna2 suppressed the generation of the aberrant products (W and Low MW) in Pif1-containing reactions (Fig. 4B, lanes 2 and 3). Dna2 is essential when Pif1 is present, but not in cells lacking Pif1^30^, suggesting that Dna2 suppresses some deleterious activity of Pif1. We suggest that the aberrant products produced by Pif1 and suppressed by Dna2 may reflect this deleterious activity. Our results indicate that Pif1, together with the replisome, can displace MCM DHs from DNA. Supp. Fig. 4E shows that Pif1 cannot remove MCM DHs from DNA in the absence of the replisome, indicating that MCM DH removal requires replisome progression. Pif1 also promoted termination and restored normal RIs on chromatin templates in the presence of excess double hexamers (Fig. 4C).

Hypomorphic *mcm10* mutants also exhibit exaggerated pausing at a dormant origin^16^. Because Mcm10 is required for initiation, we cannot omit it from reactions; however, Supp. Fig. 4D,E shows that adding more Mcm10, up to 100 times the amount required for initiation, did not promote termination in the presence of excess MCM compared to addition of Pif1.

## Discussion

The licensing factor model was conceived by Laskey and colleagues to explain how DNA lacking eukaryotic replication origins could replicate efficiently in *Xenopus* egg extracts in a manner which was nonetheless still strictly limited to once per S phase^13, 14, 15^. The model proposed a positive-acting ‘license’ was deposited randomly onto chromatin before DNA replication. Although not explicitly stated, the license was inferred to be involved in initiating replication. Random deposition presented a theoretical problem: How does the cell deal with the fact that, by chance, the distance between adjacent licenses might be so long that completion of replication would not occur before mitotic entry? To address this, they proposed that the license was deposited in excess on DNA and was erased from DNA during replication. This would ensure there were no long tracts without license and also provided a way the cell could distinguish replicated from unreplicated DNA during S phase.

We now know that the license is the MCM double hexamer which is, indeed, deposited on chromatin in excess. In yeast, deposition is primarily at specific sequences; in metazoans it is likely to be much more random. The cell cycle regulated loading of the MCM DH by ORC, Cdc6 and Cdt1 has been well-documented: it can only happen during G1 phase because the process is inhibited by cyclin dependent kinase and, in metazoans, requires the activity of the anaphase promoting complex/cyclosome. The MCM DH is the precursor of the CMG replicative helicase, so the act of initiation removes the license from the fired origin. The final piece of this model was to explain how the excess license is removed during S phase. Our results show that the complete eukaryotic replisome cannot remove the MCM double hexamer from chromatin by itself. Instead, the replisome, like the CMG by itself^11^, appears only able to push mobile MCM DHs ahead of it, presumably in much the same way CMG and RNA polymerase can push the MCM DH^11, 12^. This can continue until the MCM DH encounters some immovable object. On circular plasmids, the MCM DH can migrate until it meets a converging replication fork (Fig. 4D). At this point, the replisome is prevented from progressing through the last turns of unreplicated DNA by the MCM DH. In chromatin, nucleosomes appear to block progression of the CMG-MCM DH. CMG can progress through nucleosomes with the help of FACT/Nhp6^18^. Although we do not know how this works, FACT associates with components of the replisome^31, 32^. It is likely that the presence of the MCM DH ahead of CMG prevents FACT from accessing the nucleosome ahead of the fork.

Instead, our work indicates that an additional factor is required to remove unfired MCM DHs. We have found that the Pif1 helicase, together with the replisome, can very efficiently support MCM DH removal from both naked DNA and chromatin. The mechanism by which Pif1 acts with the replisome to remove double hexamers is unclear. Pif1 and Rrm3 have the opposite translocation polarity (5’—>3’) of the CMG (3’—>5’); concomitant action of the two helicases pushing MCM DH from the lagging and leading strands respectively may change the geometry of the two strands relative to the double hexamer, allowing the single strands to be pulled through intersubunit interfaces (Fig. 4E). It is likely that Pif1’s relative Rrm3 is most important for this function *in vivo* because *rrm3Δ* mutants show extended pausing of replication forks at dormant replication origins. Based on our results, we think it is likely that this pausing is caused by MCM DHs rather than other features of the origin like ORC or Abf1 binding. That *RRM3* is not essential suggests that there must be back up mechanisms.

Certainly Pif1 is one likely backup, and *rrm3Δ pif1-m2* double mutants are almost inviable in our hands. It is possible that other mechanisms may contribute in the absence of Rrm3 and Pif1. CMG is removed from chromatin at the end of replication via a ubiquitin- and Cdc48-mediated pathway^33, 34, 35^. This may also contribute to removing inactive double hexamers *in vivo* though the geometry and topology of CMG colliding with a double hexamer is very different from two CMGs converging at termination. It may also be that inactive double hexamers at stalled replisomes are ultimately activated and converted to active CMG, allowing normal termination and CMG removal.

Previous work showed that *mcm10* hypomorphic mutants exhibit enhanced fork pausing at a dormant replication origin^16^, suggesting that Mcm10 may have a role in deactivating dormant origins. Our results show that Mcm10 cannot promote MCM double hexamer removal without Pif1, even at levels far above those required for replication initiation. However, we cannot rule out the possibility that Mcm10 may contribute to the Pif1-dependent removal of double hexamers. Alternatively, the pause seen in *mcm10* mutants may be due to previously described defects in elongation^36, 37^, which may be exacerbated by collision with the MCM double hexamer. The completely reconstituted system described here will be of great use in understanding how DNA replication interfaces with other genomic processes like transcription, post-replication repair, sister chromatid cohesion and nucleosome inheritance.

## Acknowledgements

We are grateful to the Fermentation facility (Structural Biology STP) for growing yeast cultures and the EM STP for access to electron microscopy. This work was supported by the Francis Crick Institute, which receives its core funding from Cancer Research UK (FC001065 and FC001066), the UK Medical Research Council (FC001065 and FC001066), and the Wellcome Trust (FC001065 and FC001066). This work was also funded by a Wellcome Trust Senior Investigator Award (106252/Z/14/Z) and a European Research Council Advanced Grant (669424-CHROMOREP) to J.F.X.D and Consolidator Grant (820102-CRYOREP) to A.C.

## Materials and Methods

### Chromatin template preparation

Chromatin templates were assembled with purified Nap1, ISW1a, and histone proteins as described previously^18^.

### Molecular weight markers

18 µg HindIII-digested lambda DNA (NEB N3012S) was de-phosphorylated with 2 μL of Calf Intestinal Phosphatase (NEB M0290S) for 1 hour with shaking (1250 rpm) at 37°C, then purified with a High Pure PCR Product Purification Kit (Roche). Following this, 8 μL of the de-phosphorylated DNA was incubated with 2 μL T4 Polynucleotide Kinase (NEB M0203S), 3 μL γ^32^P-dATP (Perkin Elmer), and 5 μL H_2_O for 1 hour with shaking (1250 rpm) at 37°C. Unincorporated nucleotides were removed by passing the sample over two Illustra MicroSpin G-50 columns (GE Healthcare) and EDTA was added to a final concentration of 5 mM. The sample was stored at −20°C.

### Standard replication reactions

Reactions were carried out with shaking (1250 rpm) at 30°C. MCM loading was performed in a buffer containing 25 mM HEPES-KOH pH 7.6, 100 mM K-Glutamate, 10 mM Mg(OAc)_2_, 1 mM DTT, 0.01% NP-40-S, and 5 mM ATP. Purified ORC (10 nM), Cdc6 (50 nM), and Mcm2-7/Cdt1 (20 nM, unless otherwise stated) were added (in that order) to this mixture along with 5 nM of plasmid template DNA (pJY22). Reactions were incubated for 20 minutes (2 hours for train experiments), at which point DDK was added to 25 nM and CDK to 20 nM. The reaction was continued for a further 20 minutes and the volume increased two-fold with separate buffer and protein mixtures to give a final reaction buffer of 25 mM HEPES-KOH pH 7.5, 250 mM K-Glutamate, 10 mM Mg(OAc)_2_, 1 mM DTT, 0.01% NP-40-S, 3 mM ATP; 200 μM CTP, GTP, and UTP; 80 μM dATP, dCTP, dGTP, dTTP; and 33 nM α^32^P-dCTP. Replication was initiated by the addition of a protein master-mix containing (unless otherwise stated) 100 nM RPA, 30 nM Dpb11, 40 nM Cdc45, 20 nM Topoisomerase II, 10 nM Topoisomerase I, 20 nM Ctf4, 20 nM DNA Polymerase ε, 5 nM Mcm10, 210 nM GINS, 20 nM S-CDK, 20 nM Csm3/Tof1, 20 nM Mrc1, 20 nM RFC, 20 nM PCNA, 10 nM DNA Polymerase δ, 25 nM Sld3/7, 50 nM Sld2, and 40 nM DNA Polymerase α. For experiments requiring Okazaki fragment maturation and termination, Dna2, DNA Ligase I, and Fen1 were added to the protein master mix (or separately, as specified) to 40 nM. Where indicated, Pif1 was added to 5-10 nM. Unless otherwise stated, reactions were carried out for a further forty minutes following the addition of the protein master-mix.

### Chromatin replication reactions

For standard chromatin replication reactions, chromatin was exchanged to MCM loading buffer as described previously^18^. Replication reactions on chromatin were performed as described above, with the addition (as required) of 40 nM FACT, 400 nM Nhp6, and 50 nM INO80. These were added at the same time as the protein master mix to initiate replication. For reactions requiring an excess of MCMs loaded on chromatin, MCM loading was carried out for two hours as described previously in a volume of 20 μL. The proteins required for chromatin assembly (Nap1, ISW1a, and histone proteins) were concurrently incubated in buffer containing 10 mM HEPES-KOH pH 7.6, 50 mM KCl, 5 mM MgCl_2_, and 0.5 mM EGTA for 30 minutes on ice (in a total volume of 35.2 μL). The MCM loading reaction and the chromatin assembly proteins were then combined, along with 40 nM creatine phosphate, 3 mM ATP, and 8.4 μg creatine phosphate kinase in a total volume of 60 μL. The reaction was incubated for an additional four hours with shaking (1250 rpm) at 30°C. Prior to replication, Nap1, ISW1a, and excess histones were removed and the chromatin was exchanged to MCM loading buffer as described previously^18^.

### Post-reaction processing

For running samples under denaturing conditions, reactions were ended with 30 mM EDTA and unincorporated nucleotides were removed using Illustra MicroSpin G-50 columns (GE Healthcare). Samples were then separated on 0.6% alkaline agarose gels (30 mM NaOH and 2 mM EDTA; 15⨯ 10 cm) for 16 hours at 24 V, fixed in 5% TCA at 4°C, dried onto 3MM CHR paper (GE), and exposed to a phosphor screen (Sigma). The screens were analysed using ImageQuant software on a Typhoon Trio (GE Healthcare). For running samples under non-denaturing conditions, reactions were ended with 30 mM EDTA and incubated with 0.5% SDS and 16 μg of Proteinase K (Millipore) for 30 minutes at 37°C. Unincorporated nucleotides were then removed using Illustra MicroSpin G-50 columns (GE Healthcare) and the DNA isolated via phenol-chloroform extraction with Phase Lock Gel Tubes as per the manufacturer’s instructions (5PRIME). The DNA was purified by ethanol precipitation and resuspended in 10 μL TE pH 8. Samples requiring linearisation were then incubated with 1X Cutsmart buffer and 20 U of XhoI for 30 minutes with shaking (1250 rpm) at 37°C. The products were then separated on 0.6% native agarose gels (1X TAE; 15 x 10 cm), dried, and exposed as described above. For EtBr staining, gels were incubated in EtBr (0.5 μg/mL) for 20 minutes and imaged with an Amersham Imager 600 (GE). Replication product quantification was performed in ImageJ (National Institutes of Health).

### Two-dimensional gels

For two-dimensional gel electrophoresis, samples were separated under non-denaturing conditions in the first dimension followed by denaturing conditions in the second dimension. Prior to running second dimension, relevant lanes from the native gels were excised with a razor blade and equilibrated in alkaline running buffer (30 mM NaOH and 2 mM EDTA) for 30 minutes at 25°C. The gel slice was then laid horizontally along the top of a denaturing gel. Both dimensions were run for 20 hours at 24V in 0.6% agarose gels (15 ⨯ 10 cm).

### Loading assay

MCM loading was carried out as previously described^10^ with several modifications. ORC (10 nM), Cdc6 (50 nM), and Mcm2-7/Cdt1 (400 nM) were incubated with 5 nM of plasmid template DNA (pJY22) coupled to magnetic beads for 2 hours with shaking (1250 rpm) at 30°C. Pif1 was then added (5 nM) and the reaction was continued for a further 12 – 60 minutes. The reactions were carried out in a total volume of 40 μL in buffer containing 25 mM HEPES-KOH pH 7.6, 100 mM K-Glutamate, 10 mM Mg(OAc)_2_, 1 mM DTT, 0.01% NP-40-S, 5 mM ATP. Reactions were ended with 30 mM EDTA and the beads were washed three times in high salt buffer (45 mM HEPES-KOH pH 7.6, 0.5 M NaCl, 5 mM Mg(OAc)_2_, 1 mM EDTA, 1 mM EGTA, 10% glycerol, 0.02% NP-40-S) followed by two washes in low salt buffer (45 mM HEPES-KOH pH 7.6, 0.3 M KoAc, 5 mM Mg(OAc)_2_, 1 mM EDTA, 1 mM EGTA, 10% glycerol, 0.02% NP-40-S). Finally, beads were resuspended in 15 μL low salt buffer and the DNA was released by adding 600 units of Micrococcal nuclease (NEB M0247S) for 5 minutes with shaking (1250 rpm) at 30°C. The supernatant was then subjected to SDS-PAGE and silver staining in order to check for the presence of MCMs.

### Negative stain EM grid preparation

300-mesh copper grids with a continuous carbon film (EM Resolutions, C300Cu100) were glow-discharged for 30 s at 45 mA with a 100x glow discharger (EMS). 4-µl samples were applied to glow-discharged grids and incubated for 1 minute. Following blotting of excess sample, grids were stained by stirring in four 75-µl drops of 2% uranyl acetate for 5, 10, 15 and 20 s respectively. Excess stain was subsequently blotted dry.

### Electron microscopy data collection

Data were acquired on a FEI Tecnai LaB6 G2 Spirit electron microscope operated at 120kV and equipped with a 2K x 2K GATAN UltraScan 1000 CCD camera. 772 micrographs were collected at x21,000 nominal magnification (4.92 Å pixel size) with a defocus range of −0.5 to −1.5 µm.

### Image processing

Train termini and train middle sections were picked manually while remaining non-train particles were picked semi-automatically using e2boxer in EMAN2 v2.07. Contrast transfer function parameters were estimated by Gctf. All further image processing was performed in RELION v2.1. Train terminus particles were extracted with a box size of 128 pixels for reference-free 2D classification with CTF correction using the additional argument -- only_flip_phases. Initial 2D classification was performed with a 350 Å mask to focus classification on the terminal protein complex. Particles with terminal CMG complexes were subsequently 2D classified using a 500 Å mask to include neighbouring MCM complexes. Particles of one 2D class were further processed by particle re-extraction using a 1,260 Å box size followed by 2D classification without alignment to visualise flexibility of the MCM trains. Train middle section particles and non-train particles were extracted with a box size of 98 pixels for independent reference-free 2D classification using the same argument for CTF-correction and a mask of 350 Å or 300 Å respectively.

### Protein expression and purification

ORC, Cdc6, Mcm2-7/Cdt1, DDK, GINS, Cdc45, Dpb11, Sld2, Sld3/7, RPA, S-CDK, Topo I, Topo II, Pol ε, Pol α, Pol δ, RFC, PCNA, Csm3/Tof1, Mrc1, Ctf4, ISW1a, Nap1, Nhp6, INO80, and yeast histone proteins were expressed and purified as previously described^18, 19, 20^. Dna2, Fen1, DNA Ligase I, and FACT were expressed in *S. cerevisiae* (see strain details in supplementary tables 1-2). Cells were grown at 30°C in YP + 2% raffinose to 2-4 ⨯ 10^7^ cells per mL. Protein expression was induced by addition of galactose to 2% and growth continued for a further 3 hours at 30°C. Cells were harvested via centrifugation and resuspended in lysis buffer (see individual protocols for details), frozen drop-wise in liquid nitrogen, and crushed in a freezer mill (Spex SamplePrep 6775). Cell powder was stored at −80°C and all subsequent purification steps were carried out at 4°C.

### Dna2 purification

Cell powder from yAE49 was thawed and diluted 2:1 in 25 mM Tris-HCl (pH 7.5), 10% Glycerol, 0.5 mM βME, 150 mM NaCl (Buffer A + 150 mM NaCl) + protease inhibitors (0.5 mM AEBSF, 1 mM Leupeptin, and 10 μg/mL Pepstatin A). Cell debris was cleared by centrifugation (235,000 g, 4°C, 1 hour) and CaCl_2_ was added to 2 mM. Calmodulin Affinity Resin (Agilent Technologies) was added to the soluble extract and the sample incubated for 90 minutes at 4°C. The resin was collected in a 20 mL disposable column (Bio-Rad) and washed with 100 mL Buffer A + 150 mM NaCl + 1 mM βME + 2mM CaCl_2_. Dna2 was eluted with Buffer A + 150 mM NaCl + 1 mM βME + 2 mM EGTA + 2mM EDTA. The eluate was diluted in Buffer A + 1 mM βME so that the final concentration of NaCl was 100 mM and applied to a 1 mL Heparin HP column (GE) equilibrated in Buffer A + 1 mM βME + 100 mM NaCl. The protein was eluted over a 30 mL gradient to Buffer A + 1 mM βME + 600 mM NaCl. Peak fractions were pooled, diluted to 100 mM NaCl in Buffer A + 1 mM βME, and applied to a 1 mL MonoQ (GE) equilibrated in Buffer A + 1 mM βME + 100 mM NaCl. Dna2 was eluted over a 30 mL gradient to Buffer A + 1 mM βME + 600 mM NaCl. Peak fractions were pooled and dialysed in Buffer A + 150 mM NaCl + 5 mM βME.

### Fen1 purification

yJY31 cell powder was thawed and diluted two-fold in 25 mM Tris-HCl (pH 7.5), 10% Glycerol, 0.01% NP-40, 150 mM NaCl (Buffer B + 150 mM NaCl) + protease inhibitors (0.5 mM AEBSF, 1 mM Leupeptin, and 10 μg/mL Pepstatin A). Cell debris was cleared by centrifugation (235,000 g, 4°C, 1 hour) and FLAG M2 Affinity gel (Sigma) was added to the soluble extract. The sample was incubated for 90 minutes at 4°C before the resin was collected in a 20 mL disposable column (Bio-Rad) and washed with 100 mL Buffer B + 150 mM NaCl. Fen1 was eluted by incubating the resin with Buffer B + 150 mM NaCl + 0.5 mg/mL 3XFLAG peptide for 30 minutes, followed by a further 10 minute incubation with Buffer B + 150 mM NaCl + 0.25 mg/mL 3XFLAG peptide. The eluate was applied to a 1 mL Heparin HP column (GE) equilibrated in Buffer B + 150 mM NaCl and the protein was eluted over a 30 mL gradient to Buffer B + 1 M NaCl. Peak fractions were pooled and dialysed in Buffer B + 100 mM NaCl.

### DNA Ligase I purification

yJY33 cell powder was thawed and diluted 2:1 in 50 mM Tris-HCl (pH 7.5), 10% Glycerol, 1 mM EDTA, 10 mM βME, 400 mM NaCl (Buffer C + 400 mM NaCl) + protease inhibitors (0.5 mM AEBSF, 1 mM Leupeptin, and 10 μg/mL Pepstatin A). Cell debris was cleared by centrifugation (235,000 g, 4°C, 1 hour) and FLAG M2 Affinity gel (Sigma) was added to the supernatant. The sample was incubated for 90 minutes at 4°C before the resin was collected in a 20 mL disposable column (Bio-Rad) and washed with 100 mL Buffer C + 400 mM NaCl. DNA Ligase I was eluted by incubating the resin with Buffer C + 400 mM NaCl + 0.5 mg/mL 3XFLAG peptide for 30 minutes, followed by a further 10 minute incubation with Buffer C + 400 mM NaCl + 0.25 mg/mL 3XFLAG peptide. The eluate was applied to a 1 mL MonoQ column (GE) equilibrated in Buffer C + 100 mM NaCl and the protein was eluted over a 30 mL gradient to Buffer C + 750 mM NaCl. Peak fractions were pooled and dialysed in Buffer C + 100 mM NaCl.

### FACT purification

Cell powder from yAE90 was thawed and diluted two-fold in 20 mM Tris-HCl (pH 8.0), 10% Glycerol, 500 mM NaCl (Buffer D) + protease inhibitors (0.5 mM AEBSF, 1 mM Leupeptin, and 10 μg/mL Pepstatin A). Cell debris was cleared by centrifugation (235,000 g, 4°C, 1 hour) and FLAG M2 Affinity gel (Sigma) was added to the soluble extract. The sample was incubated for 90 minutes at 4°C before the resin was collected in a 20 mL disposable column (Bio-Rad) and washed with 100 mL Buffer D. FACT was eluted by incubating the resin with Buffer D + 0.5 mg/mL 3XFLAG peptide for 30 minutes, followed by a further 10 minute incubation with Buffer D + 0.25 mg/mL 3XFLAG peptide. The eluate was pooled and applied to a Superdex S200 column as previously described (Kurat et al. 2017).

### Pif1 purification

The nuclear isoform of Pif1 (amino acids 40-859) was cloned into the pET28a (Novagen) expression vector to add an amino-terminal 6XHis tag, then transformed into BL21 Rosetta cells (Merck). Cells were grown in LB at 37°C to an A_600_ of 0.5 and protein expression was induced with 0.1 mM IPTG for 20 hours at 16°C. Cells were then harvested by centrifugation and the pellet was resuspended in 40 mM KXPO_4_ pH 7.0, 100 mM NaCl, 8 mM MgCl_2_, 0.01% NP-40-S, 1 mM DTT, 10% glycerol (buffer E + 100 mM NaCl + 1 mM DTT) + protease inhibitors (0.5 mM AEBSF, 1 mM Leupeptin, and 10 μg/mL Pepstatin A).

Cells were lysed via sonication and insoluble material was cleared by centrifugation (257,000 g, 4°C, 30 minutes). The supernatant was then recovered and incubated with Ni-NTA agarose (Thermo Scientific) for 2 hours at 4°C. The resin was collected in a 20 mL disposable column (Bio-Rad) and washed extensively with buffer E + 300 mM NaCl + 1 mM DTT + 2 mM ATP + 20 mM imidazole. Pif1 was then eluted with buffer E + 150 mM NaCl + 1 mM DTT + 200 mM imidazole. The eluate was diluted two-fold in buffer E + 1 mM DTT + 150 mM NaCl and applied to a MonoS column (GE Healthcare) pre-equilibrated in buffer E + 150 mM NaCl + 1 mM DTT + 0.5 mM EDTA. Proteins were eluted over a 30 CV gradient from 150 mM NaCl to 1 M NaCl in buffer E. Peak fractions were pooled, concentrated with an Amicon Ultra 30,000 MWCO centrifugal filter (Millipore), and applied to a Superdex 200 Increase 10/300 GL column (GE Healthcare) pre-equilibrated in buffer E + 150 mM NaCl + 1 mM DTT + 0.5 mM EDTA. Following this, peak fractions were pooled, concentrated, and stored in aliquots at −80°C.

---

**Supplementary Figure 1.**
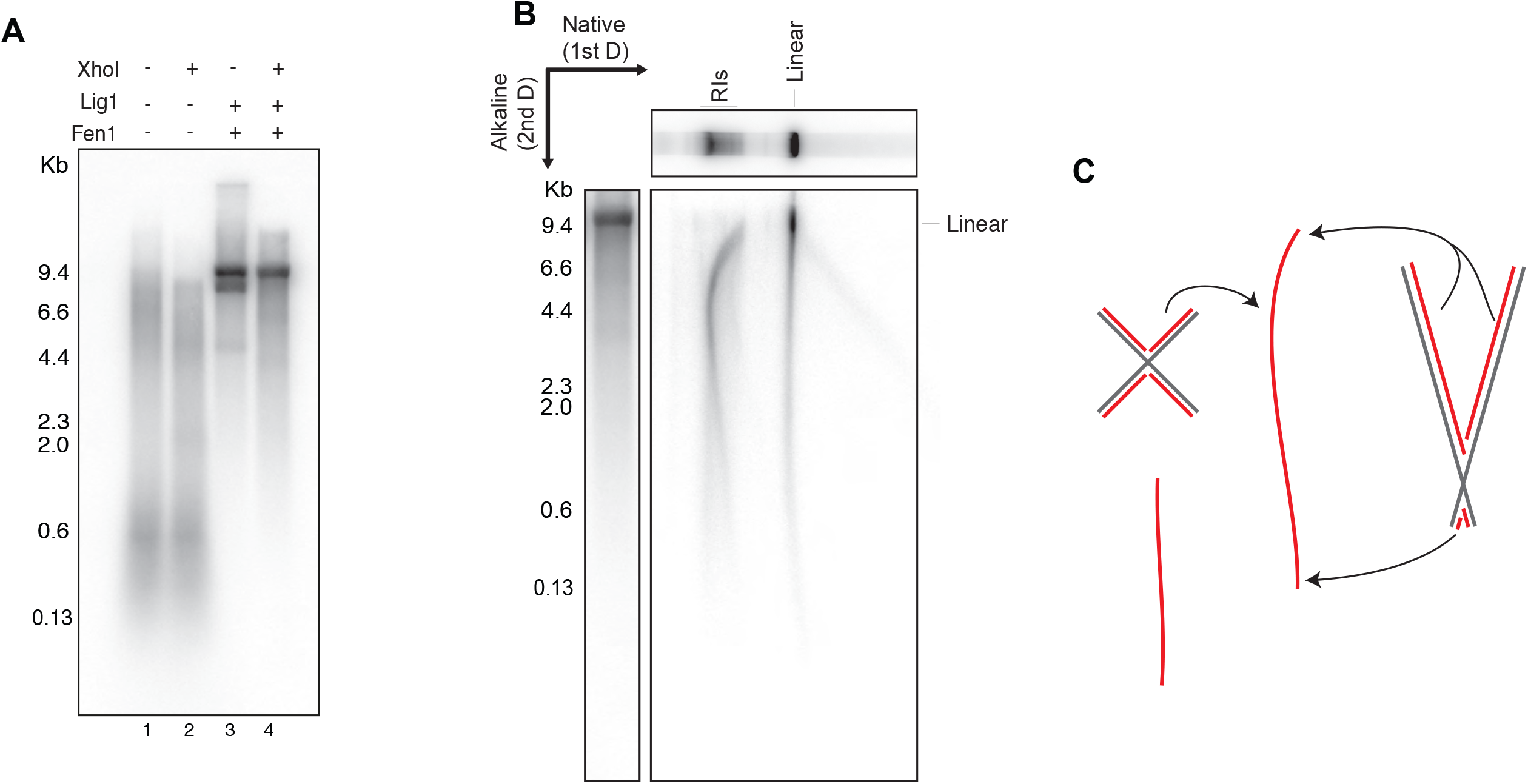
Characterisation of replication products. **a**, Reactions were performed with or without Fen1 and Lig1. Half of the products were then linearised prior to separation on an alkaline agarose gel. **b**, Two-dimensional gel analysis of linearised replication products. **c**, Schematic depicting the migration patterns of X-shaped reaction intermediates in (**b**).

**Supplementary Figure 2.**
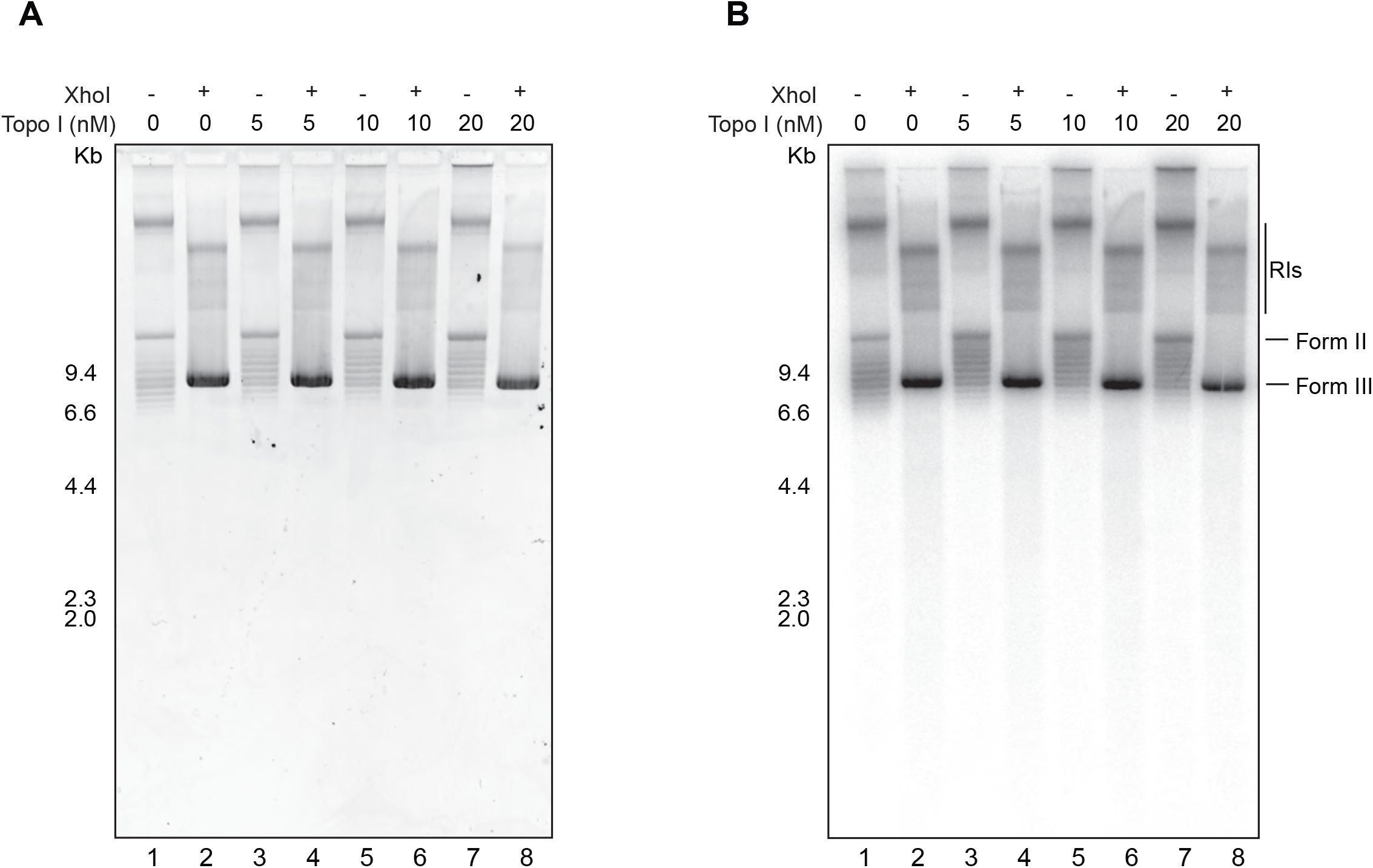
Titration of Topoisomerase I. Following replication for an hour half of the products were linearised with XhoI. They were then separated on a native agarose gel and stained with ethidium bromide (**a**) prior to drying and subjecting to autoradiography (**b**).

**Supplementary Figure 3.**
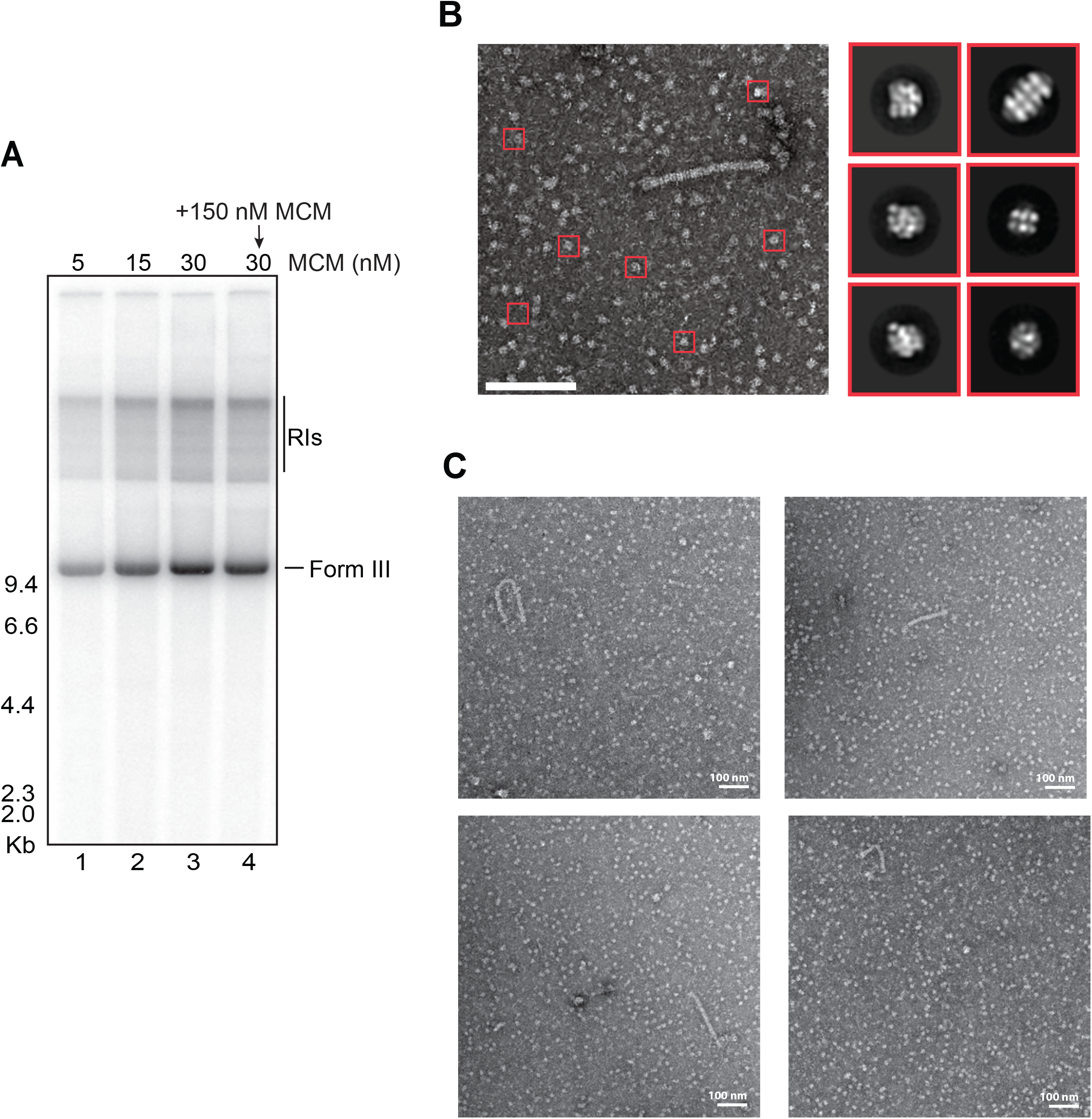
Loaded MCM double hexamers block termination. **a**, Reactions with increasing amounts of MCMs loaded onto DNA. Products were linearised and separated on a native agarose gel. Soluble MCMs were added along with the firing factors to the reaction indicated (lane 4). **b**, Representative micrograph and class averages of particles surrounding trains.**c**, Representative examples of MCM trains following replication of plasmid template DNA with an excess of loaded MCMs.

**Supplementary Figure 4.**
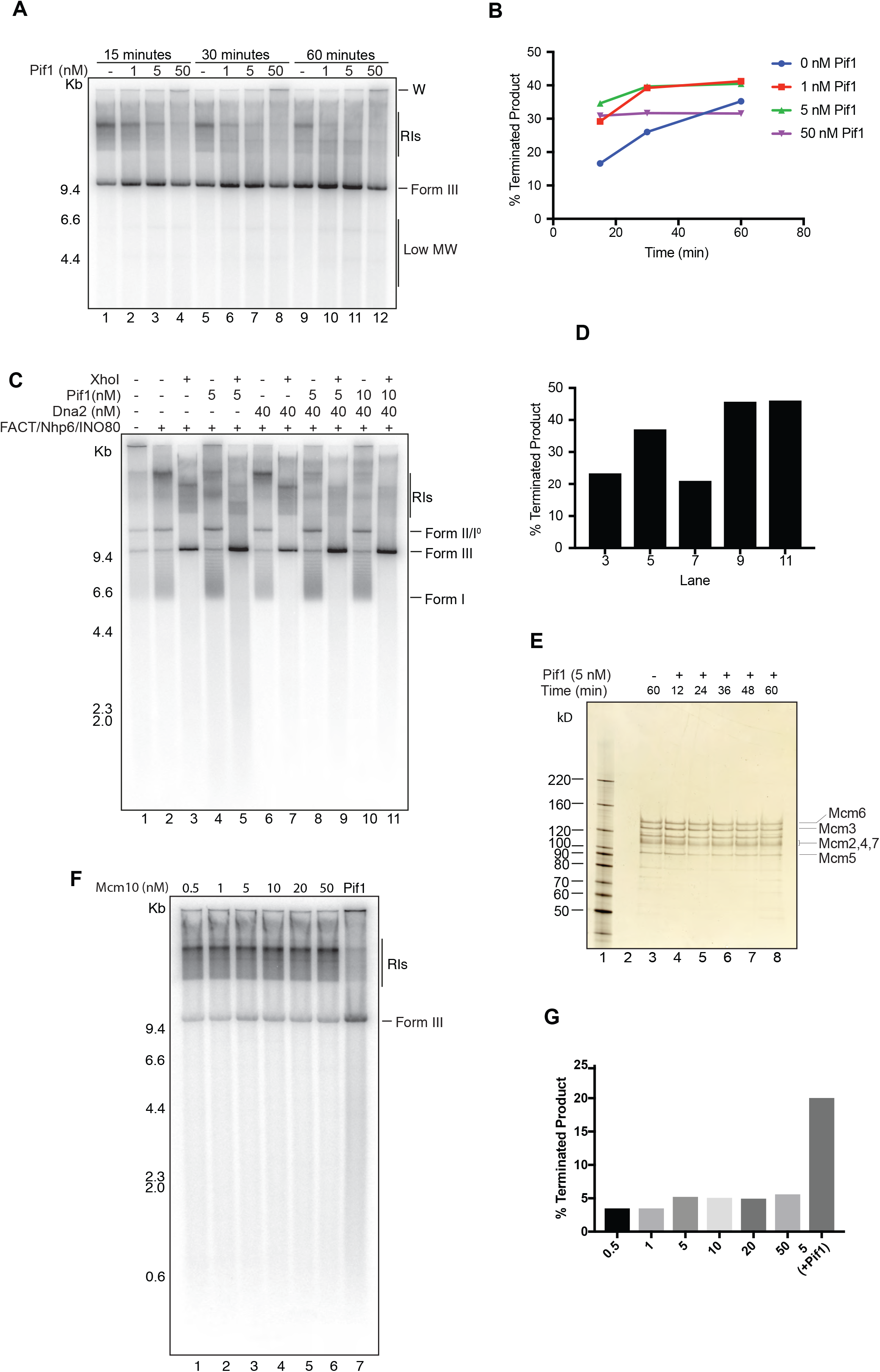
Pif1 promotes the replication-dependent removal of MCM double hexamers from DNA. **a**, Time course of replication on naked DNA performed with various concentrations of purified Pif1. Products were linearised and separated on a native agarose gel. **b**, Quantification of the terminated (linear) product as a percentage of the total product in each lane of **a. c**, Reactions performed on chromatin with various combinations and concentrations of FACT/Nhp6, Pif1, and Dna2. Products were split and half were linearised prior to separation on a native agarose gel. **d**, Quantification of the terminated product as a percentage of the total product in specified lanes of **c. e**, Loading reaction in which MCMs were loaded onto immobilised DNA and incubated for up to an hour with or without Pif1. Products were subjected to a high salt wash and analysed by SDS-PAGE and silver staining. **f**, Mcm10 titration carried out on DNA with an excess of loaded MCMs in the presence or absence of Pif1. Products were linearised and separated on a native agarose gel. **g**, Quantification of the terminated products in **f**.

**Supplementary Table 1.**
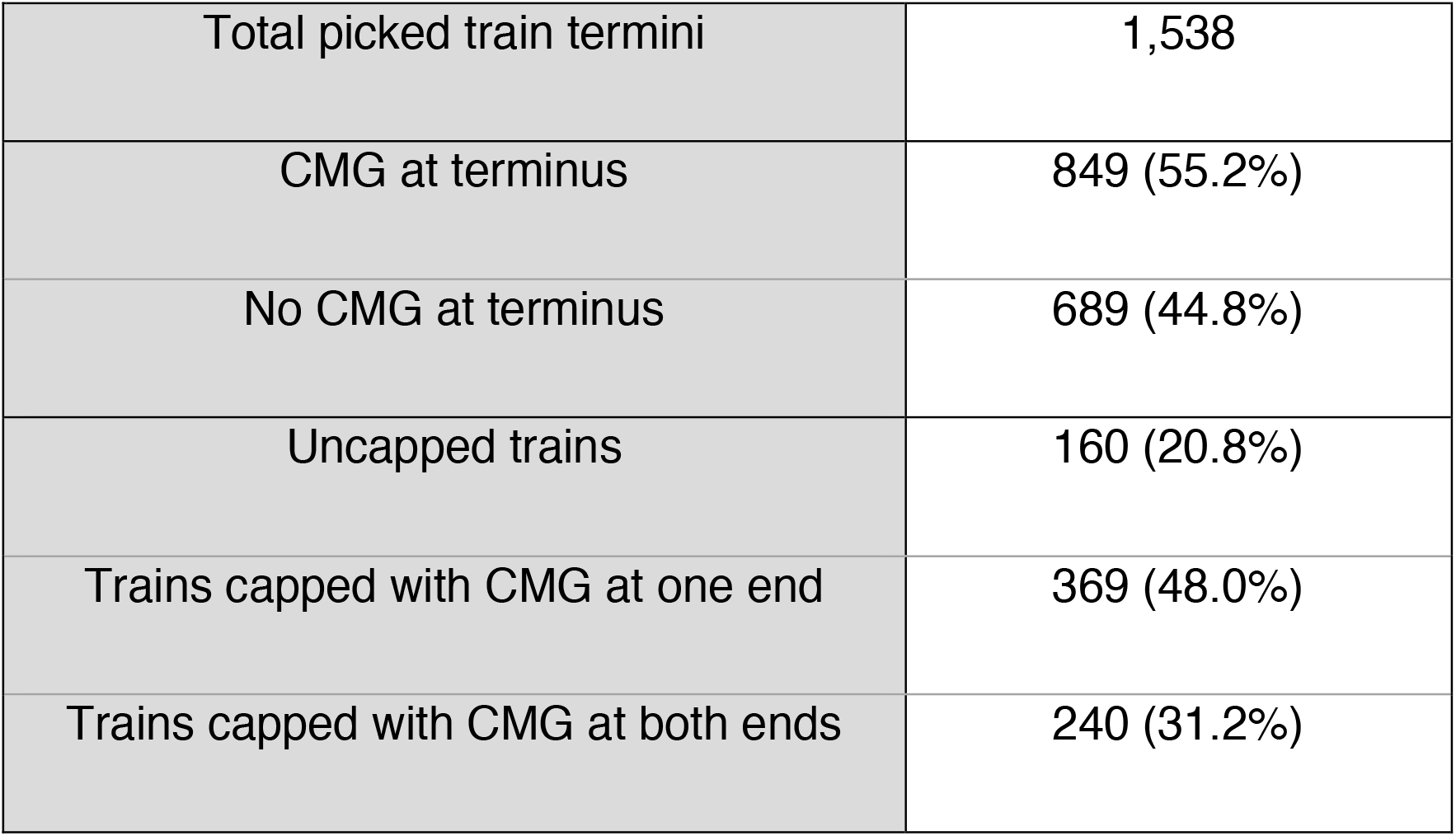
Statistical distribution of train termini.

